# Structure of mycobacterial cytochrome *bcc* in complex with Q203 and TB47, two anti-TB drug candidates

**DOI:** 10.1101/2021.06.15.448498

**Authors:** Shan Zhou, Weiwei Wang, Xiaoting Zhou, Yuying Zhang, Yuezheng Lai, Yanting Tang, Jinxu Xu, Xiaolin Yang, Luke W. Guddat, Quan Wang, Yan Gao, Zihe Rao, Hongri Gong

## Abstract

Pathogenic mycobacteria pose a sustained threat to global human health. Recently, cytochrome *bcc* complexes have gained interest as targets for antibiotic drug development. However, there is currently no structural information for the cytochrome *bcc* complex from these pathogenic mycobacteria. Here, we report the structures of *M. tuberculosis* cytochrome *bcc* alone (2.68 Å resolution) and in complex with clinical drug candidates Q203 (2.67 Å resolution) and TB47 (2.93 Å resolution) determined by single-particle cryo-electron microscopy. *M. tuberculosis* cytochrome *bcc* forms a dimeric assembly with endogenous menaquinone/menaquinol bound at the quinone/quinol binding pockets. Q203 and TB47 are bound to the quinol-binding site. Hydrogen bonds are formed between the inhibitor and the side chains of _QcrB_Thr313 and _QcrB_Glu314, residues that are conserved across pathogenic mycobacteria. These high-resolution structures provide a basis for the design of new mycobacterial cytochrome *bcc* inhibitors that could be developed into broad spectrum drugs to treat mycobacterial infections.

## Introduction

Mycobacteria, which belong to the phylum Actinobacteria, have coevolved with humans over thousands of years (Chisholm et al., 2016). Approximately 200 species of mycobacteria have been identified that have diverse lifestyles, morphologies, biochemistries and physiologies (Tortoli et al., 2017). Mycobacteria can broadly be grouped into two categories: tuberculosis-causing mycobacteria and non-tuberculous mycobacteria (NTM). *Mycobacterium leprae* is often represented in a distinct genetic clade owing to its genetic and phenotypic differences compared to other mycobacterium species (Cole et al., 2001). Mycobacteria can be further classified into fast-growing and slow-growing species or species complexes, these assignments are according to the physiological, phenotypic and phylogenetic characteristics (Rastogi et al., 2001). Although nearly all mycobacteria are saprophytes or non-pathogenic to humans, a few species cause diseases resulting in pulmonary and extra-pulmonary infections that can affect nearly all organs. Infections, which are caused by strict or opportunistic pathogenic mycobacteria (***Figure 1***) pose a sustained threat to human health. Of these, tuberculosis (TB), caused by *Mycobacterium tuberculosis* (*Mtb*), is the most serious, leading to ~ 1.2 million fatalities per year (World Health Organization, 2019). Infections involving other pathogenic mycobacteria, e.g. *M. abscessus* and *M. avium complex*, are on the rise with some outnumbering those caused by *M. tuberculosis* in countries including the United States (Donohue, 2018; Johansen et al., 2020). These infections are notoriously difficult to treat due to intrinsic or emerging resistance to many common antibiotics, thus exacerbating the challenge to find suitable drug targets.

**Figure 1.**
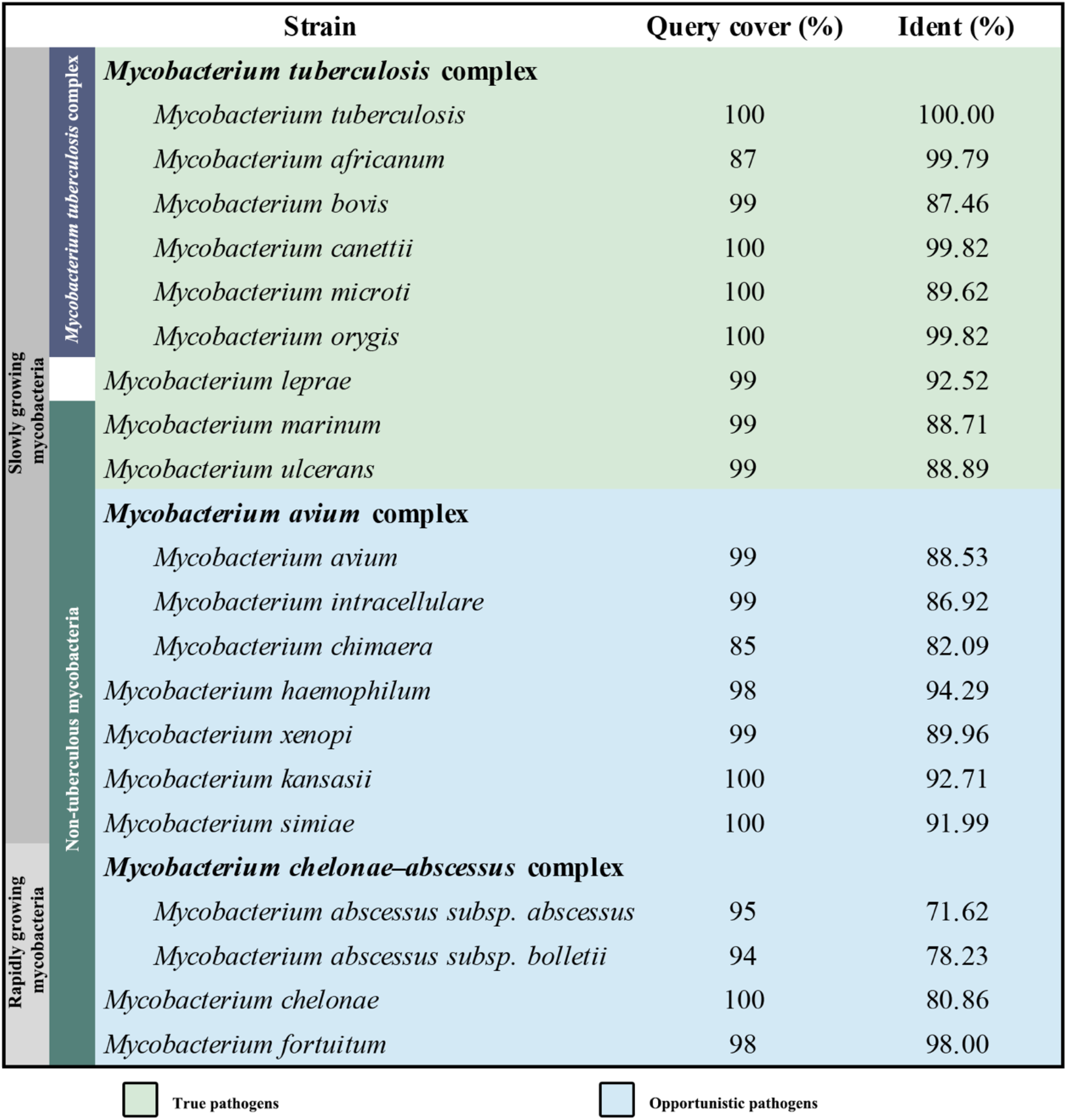
Sequences similarity comparison of *M. tuberculosis* QcrB with other pathogenic mycobacteria.

Oxidative phosphorylation (OXPHOS) has gained interest as a target space for antibiotic drug development (Cook et al., 2017; Hards et al., 2020). In OXPHOS, the protein complexes of the electron transport chain (ETC) establish a proton motive force (PMF) across a biomembrane that drives the synthesis of adenosine triphosphate (ATP) by ATP synthase (Mitchell, 1961). Maintenance of PMF and ATP homeostasis is required for the survival of both replicative and non-replicative (often referred to as dormant) mycobacteria and its dissipation leads to a rapid loss of cell viability and cell death (Koul et al., 2008; Rao et al., 2008). The reliance on the PMF and ATP homeostasis thus highlights the importance of the mycobacterial proton-pumping cytochrome *bcc-aa*_3_ supercomplex, which consists of a *bcc* menaquinol reductase (complex III, CIII) and an *aa*_3_ oxidase (complex IV, CIV) that are tightly associated (Gong et al., 2018; Kim et al., 2015; Megehee et al., 2006). Several studies support the attractiveness of cytochrome *bcc-aa_3_* for mycobacterial drug development (de Jager et al., 2020; Liu et al., 2019; Lu et al., 2018; Pethe et al., 2013; Scherr et al., 2018). Given the strict sequence conservation of this complex (***Figure 1***), broad spectrum activity of a therapeutic within the pathogenic mycobacteria is likely (Lee et al., 2020). Interestingly, all the cytochrome *bcc*-*aa*_3_ inhibitors published to date appear to target the QcrB subunit (***Figure 1***) of the cytochrome *bcc* complex and are likely bound to the menaquinol binding (Qp) site of the QcrB subunit (Lee et al., 2020). The most advanced of these are Q203 and TB47, which have been shown to clear infections due to *M. tuberculosis* (de Jager et al., 2020; Lu et al., 2018; Pethe et al., 2013) and *M. ulcerans* (Liu et al., 2019; Scherr et al., 2018). Q203 has recently completed phase II clinical trials for TB treatment (ClinicalTrials.gov number, NCT03563599) (de Jager et al., 2020). TB47 has also been evaluated in a preclinical study (http://www.newtbdrugs.org/pipeline/discovery).

To progress an understanding of the cytochrome *bcc* structure and its interaction with new drug leads, here we have determined the atomic resolution cryo-electron microscopy (cryo-EM) structures of *M*. *tuberculosis* cytochrome *bcc* alone (2.68 Å resolution) and in complex with Q203 (2.67 Å resolution) and TB47 (2.93 Å resolution). The high resolution *M*. *tuberculosis* cytochrome *bcc* structures will greatly accelerate efforts towards structure-guided drug discovery for pathogenic mycobacteria including *M. tuberculosis*.

## Results and Discussion

### Structure of *M. tuberculosis* cytochrome *bcc*

A hybrid supercomplex consisting of *M. tuberculosis* CIII and *Mycobacterium smegmatis* CIV was purified and its structure determined by cryo-EM to an overall resolution of 2.68 Å (***Figure 2-figure supplement 1; Supplementary file 1***). The *M. tuberculosis* cytochrome *bcc* is dimeric similar to the *M. smegmatis bcc* complex in the CIII/CIV supercomplex (Gong et al., 2018) and the bovine mitochondrial cytochrome *bc_1_* complex (Iwata et al., 1998) (***Figure 2A, B***). The dimensions of the complex are 140 Å by 70 Å by 100 Å (***Figure 2A, B***). Three canonical subunits QcrA, QcrB and QcrC with all the prosthetic groups and endogenous menaquinones were clearly assigned in the unambiguous cryo-EM density (***Figure 2C; Figure 2-figure supplement 2***). QcrA has three transmembrane helices (TMHs) and has a “U” shaped structure (***Figures 2D, 3A***). The N-terminal region with TMH1/2 and the TMH3 make up the two arms of the “U” structure. These arms are linked by the region located near the cytoplasmic side. Attached to _QcrA_TMH3 is the C-terminal domain, which faces the periplasmic side of the membrane and holds the [2Fe-2S] cluster in place. QcrB has eight TMHs (***Figures 2D, 3B***). Four of these are responsible for burying two functionally important heme *b* cofactors (high potential heme *b*_H_ and low potential heme *b*_L_). Both heme *b*_L_ and heme *b*_H_ are bound between TMH I/II and TMH III/IV, heme *b*_L_ is near the periplasmic side and heme *b*_H_ near the cytoplasmic side. QcrC is a transmembrane protein with a C-terminal TMH located between _QcrB_TMH5 and _QcrB_TMH7 (***Figures 2D, 3C***). The N-terminal periplasmic portion of QcrC can be divided into two heme-containing cytochrome *c* domains designated D1 and D2. The D1 domain protrudes out of the core of CIII whereas the D2 domain interacts extensively with QcrA and QcrB.

**Figure 2.**
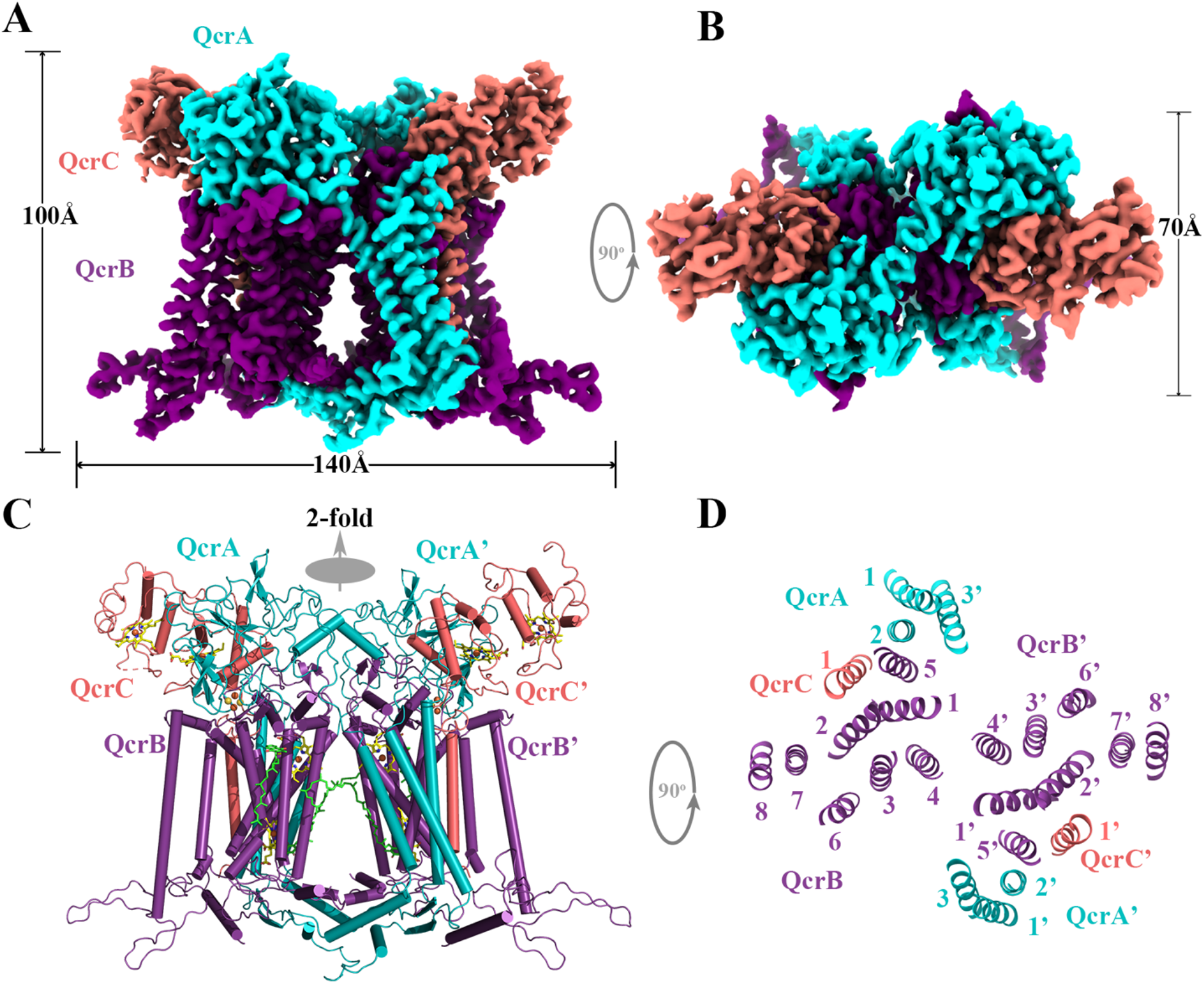
Overall architecture of the of *Mtb* cytochrome *bcc* complex. (**A**) Front view and (**B**) top view of cryo-EM map of cytochrome *bcc* at 2.68 Å resolution. QcrA, QcrB, and QcrC are colored teal, purple, and shalmon, respectively. (**C**) Cartoon representation of cytochrome *bcc*, using the same color scheme as above. The twofold symmetry of the dimer is depicted by the grey axis. The heme groups (*b*_H_, *b*_L_, *c*_D1_, and *c*_D2_) and menaquinone/menaquinol (MK_P_/MK_N_) are shown as stick models. The [2Fe-2S] clusters are shown as spheres. (**D**) A cross-sectional view (top) of cytochrome *bcc* dimer.

**Figure 3.**
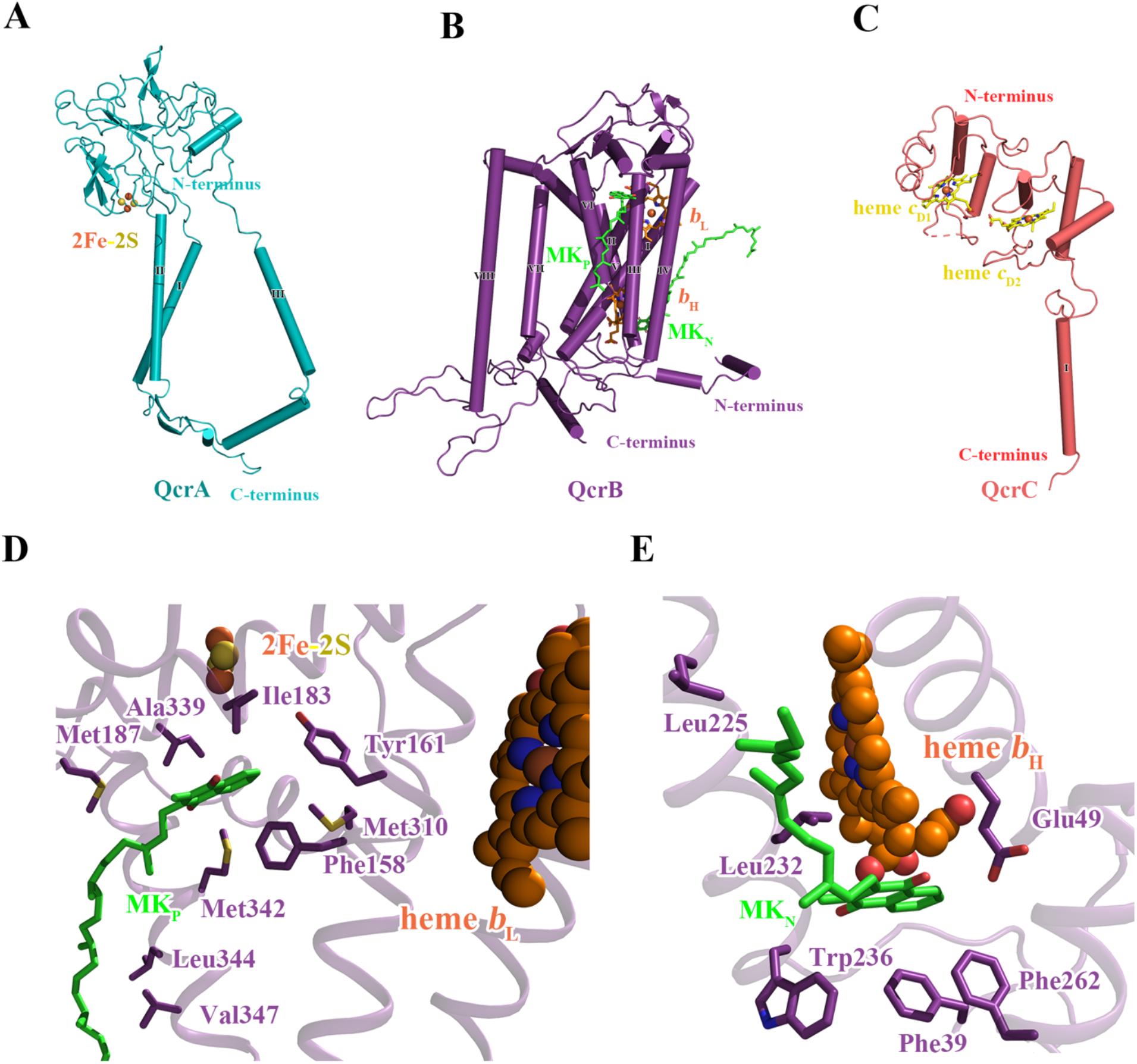
Structure of *Mtb* cytochrome *bcc* subunits. Cartoon representation of the monomers of (**A**) QcrA, (**B**) QcrB, and (**C**) QcrC, with prosthetic groups. (**D**) The Q_P_ binding site and (**E**) Q_N_ binding site. The residues potentially involved in the binding of MK/MKH_2_ are shown with side chains in a stick representation. The MK/MKH_2_ molecules are colored in green and shown as sticks. The [2Fe-2S] and heme groups are shown as spheres and labeled accordingly.

### Quinone and quinone binding pockets of *M. tuberculosis* cytochrome *bcc*

Quinone binding sites are often species-dependent and thus are important for drug discovery (Harikishore et al., 2020; Lee et al., 2020). Two quinone-binding sites could be identified (***Figure 3D, E***), the quinol oxidation site (Q_P_ site) and the quinone reduction site (Q_N_ site). The Q_P_ site responsible for menaquinol (MKH_2_) oxidation is near heme *b*_L_, whereas the Q_N_ site responsible for menaquinone (MK) reduction is close to heme *b*_H_ (***Figure 3D, E; Figure 2-figure supplement 2***). The Q_P_ site is at the center of an inverted triangle structure surrounded by helices (***Figure 3D***). One MK molecule was identified at this site with its naphthoquinone ring surrounded mainly by hydrophobic residues, _QcrB_Phe158, _QcrB_Tyr161, _QcrB_Leu180, _QcrB_Ile183, _QcrB_Met310 and _QcrB_Met342. Its hydrophobic tail that contains multiple isoprenoid groups wraps around _QcrB_TMH6 down to its cytoplasmic end by interacting with _QcrB_Met187, _QcrB_Ala339, _QcrB_Leu344 and _QcrB_Val347. However, the edge-to-edge distance from MK to heme *b*_L_ is 15 Å. Thus, we speculate that the endogenous electron donor MKH_2_ should be closer to the heme *b*_L_ to facilitate electron transfer. We speculate that what is observed here is the oxidized product as it leaves the Q_P_ site. It is worth noting that all the reported inhibitors including Q203 (Pethe et al., 2013) and TB47 (Lu et al., 2018) are suggested to interact with this Q_P_ site. In addition, the Q_N_ site is mainly formed by _QcrB_TMH1, _QcrB_TMH4, _QcrB_TMH5 and one loop region of QcrB (***Figure 3E***). The head group of MK is bound in this pocket interacting with _QcrB_Phe39, _QcrB_Glu49, _QcrB_Leu225, _QcrB_Leu232, _QcrB_Trp236 and _QcrB_Phe262, and its long hydrophobic tail extends along _QcrB_TMH1 towards the periplasmic side. MK/MKH_2_ are part of the Q-cycle hypothesis and essential for electron transfer in the cytochrome *bcc* complex (Gong et al., 2018). In addition, the electron transfer pathway of *M. tuberculosis* cytochrome *bcc* is believed to be same as that of *M. smegmatis* cytochrome *bcc* based on their highly similar cofactor arrangement (Gong et al., 2018). Given the crossspecies activity of this complex (Lee et al., 2020) and high homology of the QcrB subunits across mycobacterial pathogens (***Figure 1***), this data opens the way for the discovery of broad spectrum mycobacterial agents based on rational, structure-based inhibitor design principles.

### Q203 interactions in *M. tuberculosis* cytochrome *bcc* binding pocket

Q203 has recently been subjected to a phase II clinical study for *M. tuberculosis* treatment (de Jager et al., 2020). This compound has also been shown to be strongly bactericidal against *Mycobacterium ulcerans* (Scherr et al., 2018). It is suggested to be an inhibitor that competes with endogenous substrate binding (Q_P_ site) of the cytochrome *bcc* complex (Pethe et al., 2013), but this hypothesis is yet to be verified by direct experimental evidence. To obtain atomic information on the mode of binding of Q203 to cytochrome *bcc*, we have determined the structure of a hybrid supercomplex as described above in the presence of Q203 by cryo-EM to an overall resolution of 2.67 Å (***Figure 4-figure supplement 1 and figure supplement 2; Supplementary file 1***). The cryo-EM map shows that close to the Qp binding pocket within the membrane of each QcrB of cytochrome *bcc*, density for Q203 is present (***Figure 4A, B***). All of the Q203 molecules fill each QcrB subunit binding deeply into the Q_P_ pocket and with identical binding modes. The key interactions that anchor Q203 are (i) a hydrogen bond between the hydroxyl oxygen of the side chain of _QcrB_Thr313 and the amine in the carboxamide linker of Q203 (3.0 Å), (ii) a halogen bond between the chlorine atom of the trifluoromethyl group and an ordered water molecule that simultaneously forms a hydrogen bond with the hydroxyl oxygen of the side chain of _QcrB_Tyr164, (iii) a hydrogen bond between the side chain of _QcrB_Glu314 and the nitrogen atom in the imidazopyridine ring (3.0 Å), and (iv) a hydrogen bond between the side chain of _QcrA_His375 and the nitrogen atom in the imidazopyridine ring (2.98 Å) (***Figure 4C***). In addition, the carbon atoms of Q203 interacts with Gly^175^, Ala^179^, Leu^180^, Thr^184^, Ser^304^, Pro^306^, Met^310^, Ala^317^ and Met^342^ through hydrophobic interactions. These residues are within helices adjacent to QcrB. Consistent with these findings, functional studies have shown that substitution of _QcrB_Thr313 to alanine confers resistance to the Q203 (Pethe et al., 2013). Interestingly, the binding of Q203 involves residues from both QcrA and QcrB. Due to the need to form stabilizing interactions between subunits, resistance may be more difficult to achieve here than if the site involved only one subunit. Furthermore, the mapping of reported mutations in Q203-resistant *M. tuberculosis* reveals that they are positioned directly where Q203 binds in this structure (Lupien et al., 2020) (***Figure 4-figure supplement 3***).

**Figure 4.**
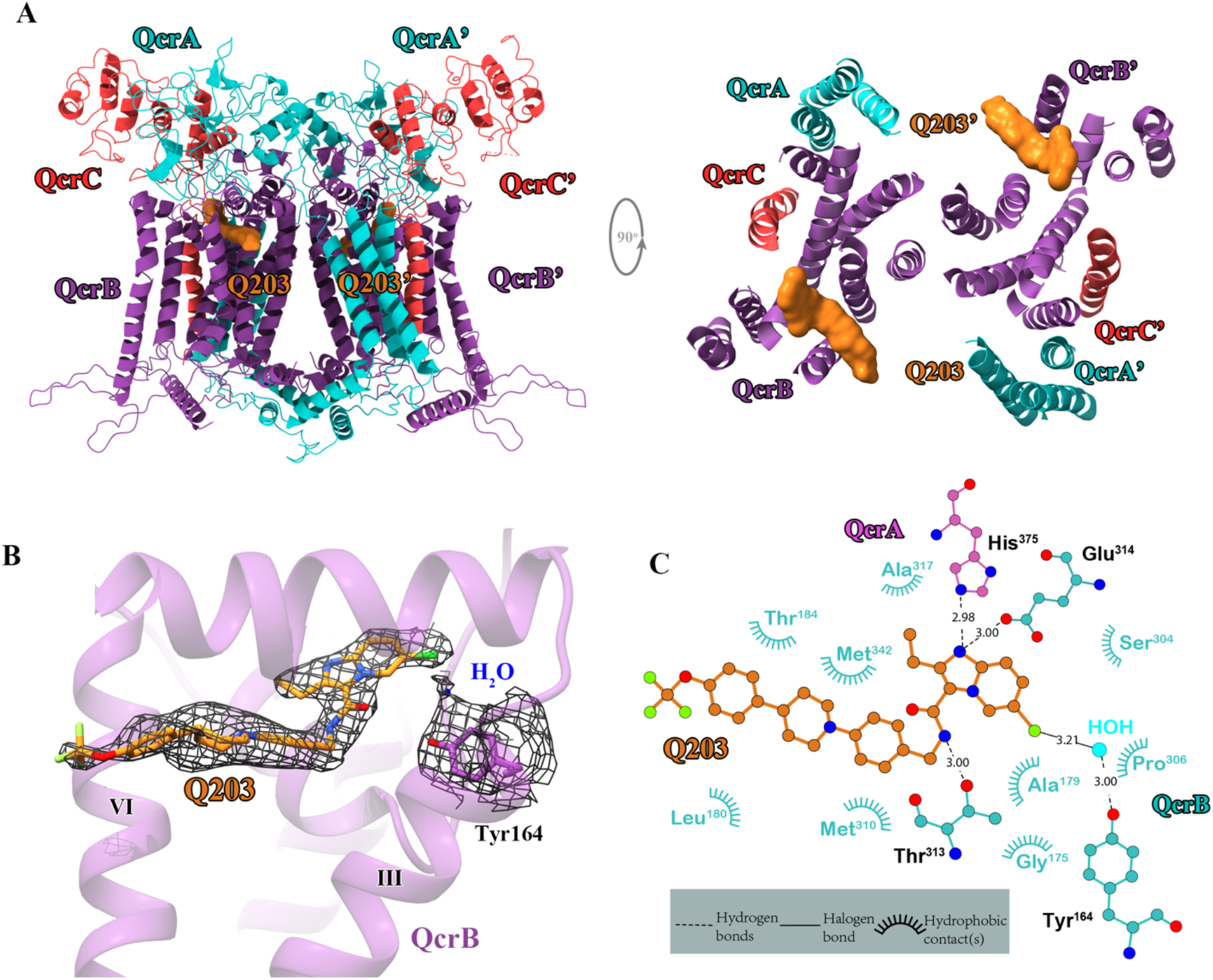
Cryo-EM structure of the *Mtb* cytochrome *bcc* complex in the presence of Q203. (**A**) Side (left) and top (right) views of the cryo-EM structure of the *Mtb* cytochrome *bcc* complex presented as a cartoon representation. Q203 (orange) is bound to the Qp site. (**B**) Visualization of densities for Q203, a water molecule and _QcrB_Tyr164. (**C**) Plot of distances of various parts of Q203 to residues in the Qp site, determined using LIGPLOT (www.ebi.ac.uk/thornton-srv/software/LIGPLOT/).

### TB47 binding mode of *M. tuberculosis* cytochrome *bcc*

TB47, currently being evaluated in preclinical studies, has also been reported to target the QcrB of cytochromes *bcc* from *M. tuberculosis* (Lu et al., 2018) and *M. ulcerans* (Liu et al., 2019). Here we have determined its structure in complex with the hybrid mycobacterial cytochrome *bcc*. The 2.93 Å cryo-EM map shows density for TB47 and confirms that it binds in the same location as Q203 (***Figure 5A, B; Figure 5-figure supplement 1 and figure supplement 2; Supplementary file 1***). Three hydrogen bond interactions are observed involving the side chains of _QcrB_Thr313, _QcrB_Glu314, and _QcrA_His375. Similar interactions are also observed when Q203 binds (***Figure 5C***). Tyr^161^, Leu^171^, Gly^175^, Ala^179^, Leu^180^, Thr^184^, Met^187^, Leu^194^, Ser^304^, Gly^305^ and Met^342^ also contribute to TB47 binding, largely through hydrophobic interactions (***Figure 5C***). A mutation in TB47-resistant *M. smegmatis (M. tuberculosis:* H195Y) is positioned close to the Qp-binding site (Lu et al., 2018) (***Figure 5-figure supplement 3***). As a result, it causes indirect steric interference with the binding of TB47, thus this structure provides the molecular basis for conferring resistance.

**Figure 5.**
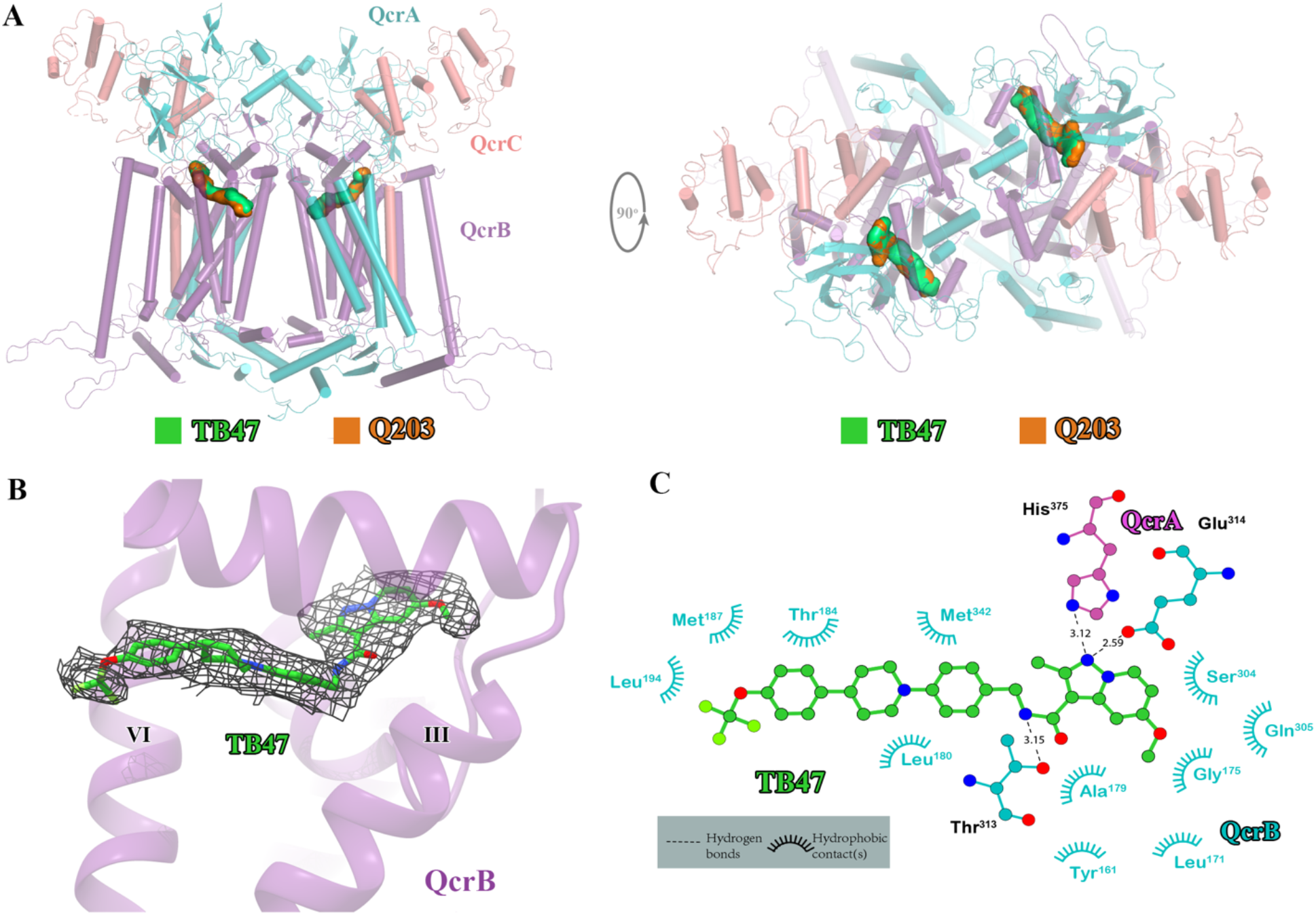
Cryo-EM structure of the *Mtb* cytochrome *bcc* complex in the presence of TB47. (**A**) Side (left) and top (right) views of the cryo-EM structure of the *Mtb* cytochrome *bcc* complex presented as a cartoon representation. The TB47 (green) and Q203 (orange) are bound to the Qp site. (**B**) Visualization of the density forTB47. (**C**) Plot of distances of various parts of TB47 to residues in the Qp site, determined using LIGPLOT (www.ebi.ac.uk/thornton-srv/software/LIGPLOT/).

### Specificity of Q203 and TB47 for mycobacterial cytochromes *bcc* complex

The basis for the high specificity of Q203 and TB47 toward the Qp site of mycobacterial cytochromes *bcc* becomes apparent in the structural comparison between the QcrB subunit of *M. tuberculosis* and counterparts from other species (***Figure 6***). The highly conserved residues that are involved in the binding of these two molecules in this region (***Figure 6-figure supplement 1***) suggest a consistent overall fold and binding site exists in mycobacteria. This is also in agreement with the fact that Q203 and TB47 show antimycobacterial activity across many species (de Jager et al., 2020; Liu et al., 2019; Lu et al., 2018; Pethe et al., 2013; Scherr et al., 2018). In contrast, in other prokaryotic, eukaryotic and human Qp-binding pockets, for example, from *Saccharomyces cerevisiae* (Lange and Hunte, 2002), *Rhodobacter sphaeroides* (Esser et al., 2008) or human (Guo et al., 2017), many of the observed interactions would be sterically hindered (***Figure 6***). This suggests that Q203 and TB47 should have low binding affinity toward its counterpart QcrB in non-mycobacterial bacteria and in eukaryotes. Coincidentally, the residues contributing to the clashes of Q203 and TB47 in the Qp binding pockets are commonly observed (***Figure 6***). Even if there is some flexibility in the Qp binding pocket that enables some level of binding, key residues that enable the binding of Q203 and TB47 in the mycobacteria are not present in other bacteria and eukaryotes (***Figure 6-figure supplement 1***). These observations correlate with low general antibacterial activity and low cytotoxicity for Q203 and TB47 (Liu et al., 2019; Lu et al., 2018; Pethe et al., 2013; Scherr et al., 2018).

**Figure 6.**
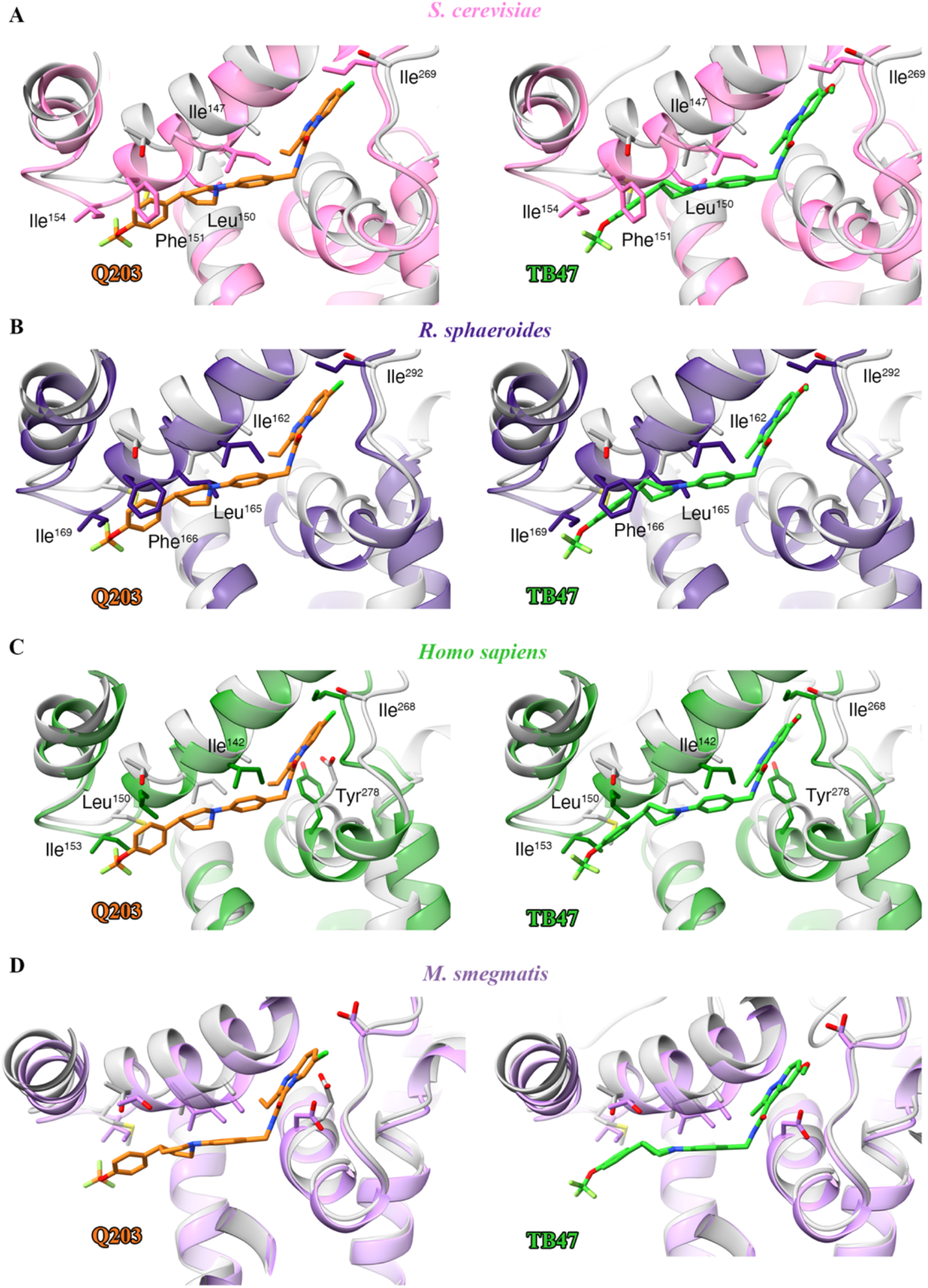
Structural alignment between the *Mtb* Qp binding pocket where Q203 or TB47 binds with homologous subunits from four other species. These subunits are from *S. cerevisiae* (pink, PDB: 1KYO), *R. sphaeroides* (blue, PDB: 2QJP), *Homo sapiens* (green; PDB: 5XTE), and *M. smegmatis* (violet, PDB: 6ADQ). Residues causing steric clashes in the homologous subunits are labeled. Q203 and TB47 are shown as orange and green stick models, respectively.

### Implication of Q203 and TB47 inhibitory mechanism

To gain further insights into the mechanism of action of Q203, we compared the structures of *M. tuberculosis* cytochrome *bcc* in the presence and absence of Q203 (***Figure 7-figure supplement 1A***). The structure of apo cytochrome *bcc* is almost identical with the Q203-bound structure (rmsd of 0.497 Å for all Ca atoms), which suggests that Q203 binding does not significantly affect the overall architecture of cytochrome *bcc*. A comparison of the Q203-bound and Q203-free cytochrome *bcc* structures shows residues involved in the binding pocket move outward, thus adapting to the shape of Q203 (***Figure 7-figure supplement 1B***). Specifically, the side chains of _QcrB_Ser304, _QcrB_Glu313, _QcrB_Glu314 and _QcrB_Met342 undergo significant conformational changes to form hydrogen bonds to Q203. The binding of TB47 to *M. tuberculosis* cytochrome *bcc* also induces very similar conformational changes in the Qp binding pocket to those seen for Q203 (rmsd of 0.454 Å for all Ca atoms) (***Figure 7-figure supplement 1B***). Differences in binding are due to the different ethyl group and methyl moieties in the head groups of Q203 and TB47, respectively. It is also important to note that one endogenous substrate molecule is also bound to the Qp site in the apo structure of cytochrome *bcc*, which potentially affects the evaluation of the conformational changes upon the binding of Q203 or TB47.

When analyzing the superimposed structures (***Figure 7-figure supplement 1***), it is apparent that Q203 and TB47 act competitively with the quinol binding as they almost completely occupy the Q_p_ pocket. We therefore conclude that Q203 and TB47 are bona fide analogs of the substrate, and thus ultimately function by hindering the downstream synthesis of ATP (***Figure 7A***). These two compounds are also highly bactericidal against *M. ulcerans*, almost certainly targeting the Qp-binding site (Liu et al., 2019; Scherr et al., 2018). In summary, the sequences of the QcrB subunits have high homology across pathogenic mycobacteria (Lee et al., 2020) and the essential residues (_QcrB_Glu313 and _QcrB_Glu314) that are involved in hydrogen-bonding interactions with the inhibitors (Pethe et al., 2013; Scherr et al., 2018) are conserved across pathogenic mycobacteria (***Figure 7B***).

**Figure 7.**
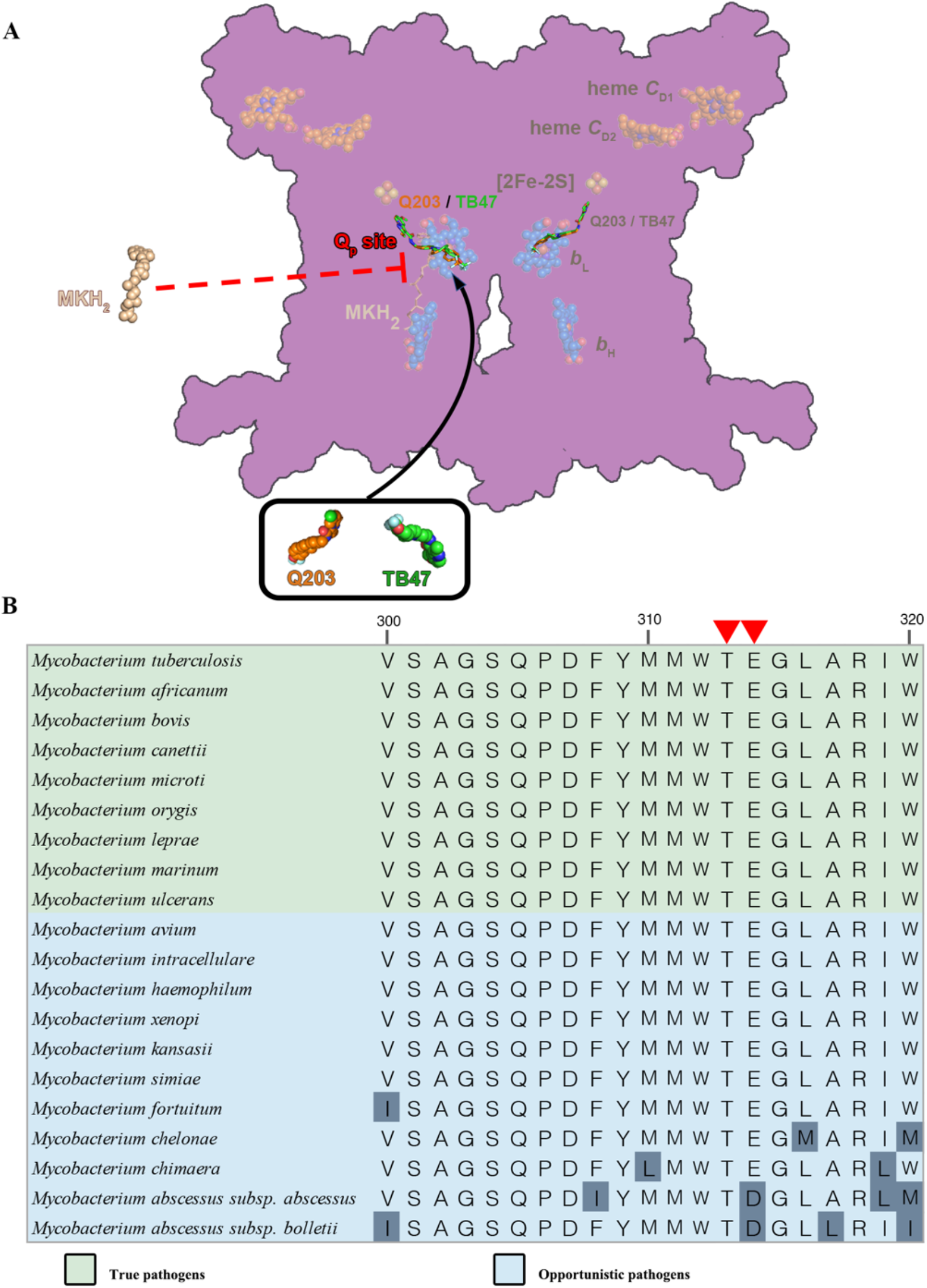
Proposed mechanism of inhibition and sequence alignment for the Q_P_ site within pathogenic mycobacteria. (**A**) Schematic of *M. tuberculosis* cytochrome *bcc* inhibition by Q203 and TB47. *M. tuberculosis* cytochrome *bcc* dimer are colored magenta. The binding of Q203 (orange spheres) or TB47 (green spheres) prevent substrate access (gray spheres). The prosthetic groups are shown in spheres and labeled accordingly. (**B**) Partial sequence alignment at the Qp site. The red triangles indicate the conservative residues involving with the hydrogen-bond formation between Q203 and TB47.

## Conclusions

We have determined the apo- and Q203 and Tb47-bound structures of a hybrid pathogenic *M. tuberculosis/M. smegmatis* cytochrome *bcc* complex. The study shows the structural features of *M. tuberculosis* cytochrome *bcc* of and how it is specifically inhibited by Q203 and TB47. The extensive interactions between Q203 or TB47 and the Qp binding pocket account for the highly specific binding of these two inhibitors to pathogenic *M. tuberculosis* cytochromes *bcc* compared to eukaryotic counterparts. Two conservative residues involved with the formation of hydrogen bonds are observed across the pathogenic mycobacteria. These structures provide a long-sought basis for rational, structure-based inhibitor design to accelerate the development of Q203 and TB47 analogs as drug leads for mycobacterial infections.

## Materials and Methods

### Key resources table

| REAGENT or RESOURCE | SOURCE | IDENTIFIER |
| --- | --- | --- |
| **Bacterial and Virus Strains** |  |  |
| *M. smegmatis* MC<sup>2</sup> 51 | (Li et al., 2014) |  |
| **Chemicals, Peptides, and Recombinant Proteins** |  |  |
| Lauryl Maltose Neopentyl Glycol (LMNG) | Anatrace | Cat# NG310 |
| Digitonin | BIOSYNTH | Cat# D-3200 |
| **Critical Commercial Assays** |  |  |
| Ni-NTA Agarose | QIAGEN | Cat# 30230 |
| Superdex 200 increase | GE Healthcare | Cat# 28990944 |
| **Deposited Data** |  |  |
| EM map apo complex | This paper | EMDB code: EMD-30943 |
| Apo complex structure | This paper | PDB code: 7E1V |
| EM map Q203-bound complex | This paper | EMDB code: EMD-30944 |
| Q203-bound complex structure | This paper | PDB code: 7E1W |
| EM map TB47-bound complex | This paper | EMDB code: EMD-30945 |
| TB47-bound complex structure | This paper | PDB code: 7E1X |
| **Software and Algorithms** |  |  |
| SerialEM | (Mastronarde, 2003) | http://bio3d.colorado.edu/SerialEM/ |
| MotionCor2 | (Zheng et al., 2017) | https://emcore.ucsf.edu/ucsf-motioncor2 |
| RELION | (Zivanov et al., 2019) | https://www2.mrc-lmb.cam.ac.uk/relion |
| cryoSPARC | (Punjani et al., 2017) | https://cryosparc.com/ |
| Phyre2 | (Kelley et al., 2015) | http://www.sbg.bio.ic.ac.uk/phyre2/html/page.cgi?id=index |
| UCSF Chimera | (Pettersen et al., 2004) | https://www.cgl.ucsf.edu/chimera/ |
| COOT | (Emsley et al., 2010) | https://www2.mrc-lmb.cam.ac.uk/personal/pemsley/coot/ |
| PHENIX | (Adams et al., 2010) | https://www.phenix- |
| PyMOL | (Schrodinger, 2010) | online.org/<br>https://pymol.org/2/ |
| **Other** |  |  |
| 100kDa cutoff concentrators | Sartorius | Cat# VS0642 |
| R0.6/1.0 300 mesh Cu holey carbon grids | Quantifoil |  |

### Expression of hybrid supercomplex consisting of *Mtb* CIII and *Msm* CIV

The hybrid supercomplex was obtained according to a previous study (Kim et al., 2015) but with some modifications. The *Mtb* cytochrome *bcc* complex is encoded by three putative genes (Rv2194-2196). Genes were amplified from H37Rv genomic DNA by PCR using Phanta Max DNA polymerase (Vazyme), and two steps PCR was used to inset a 10 × His tag at the C-terminus of the qcrB (Rv2196). Genes encoding the entire cytochrome *bcc* complex operon were then cloned into the vector pVV16. The resultant plasmid was transformed into *M. smegmatis* mc^2^ 51 (Li et al., 2014) cells whose qcrCAB operon encoding *Msm* cytochrome *bcc* had already been knocked out. The cells were cultivated in Luria-Bertain broth (LB) liquid media supplemented with 50 μg/mL hygromycin, 25 μg/mL streptomycin and 0.1% Tween 80. Cell pellets were harvested by centrifugation when the cells were grown to an optical density (OD_600_ of 1.0-1.2) at 37 °C (220rpm). Harvested cells were frozen at −80 °C until use.

### Purification of the hybrid supercomplex

Cell pellets were thawed and resuspended in buffer A containing 20 mM MOPS, pH 7.4, 100 mM NaCl, and then lysed by passing through a French Press at 1,200bar three times. Cell debris and non-lysed cells were removed by centrifugation at 14,000 rpm for 10 min at 4 °C. The supernatant was collected and ultra-centrifuged at 36,900 rpm and 4 °C for 2 h. The membrane fraction was solubilized by addition of 1% (w/v) LMNG (lauryl maltose neopentyl glycol) in buffer A and incubated for 2 h at 4 °C with slow stirring. The suspension was ultra-centrifuged and the supernatant was applied to Ni-NTA agarose beads (GE Healthcare) at 4 °C. The beads were further washed in buffer A with 50 mM imidazole and 0.004% (w/v) LMNG. The buffer was exchanged to buffer B (20 mM MOPS, pH 7.4, 100 mM NaCl and 0.1% (w/v) digitonin) and then washed in resin in batch mode. The protein was eluted from the beads with buffer B containing 500 mM imidazole. Protein was then concentrated and loaded onto a Superdex 6 increase (10/300 GL, GE Healthcare) column equilibrated in buffer B. Peak fractions were pooled and concentrated to ~ 8 mg/ml for electron microscopy studies.

### Cryo sample preparation and data collection

300-mesh Quantifoil R0.6/1.0 grids (Quantifoil, Micro Tools GmbH, Germany) were glow-discharged at H_2_/O_2_ atmosphere for 25s. 3 μL aliquots of protein complex at a concentration of 10 mg/mL were applied to the grid and then blotted for 3s with force 0 at 8°C and 100% humidity using a Vitrobot IV (Thermo). Images were collected using a Titan Krios 300keV electron microscope (Thermo), equipped with K3 Summit direct electron detector camera (Gatan). Data was recorded at 29,000× magnification with a calibrated super-resolution pixel size 0.82 Å/pixel. The exposure time was set to 2.4 s with 40 subframes and a total dose of 60 electrons per Å^2^. All images were automatically recorded using SerialEM with a defocus range from 1.2 μm to 1.8 μm (Mastronarde, 2003). For the datasets of apo, Q203-bound and TB47-bound *M. tuberculosis* cytochrome *bcc*, a total of 4,141, 3,763 and 2,968 images were collected, respectively.

### Image processing

All dose-fractioned stacks were motion-corrected and dose-weighted using MotionCorr2 (Zheng et al., 2017) in RELION (Zivanov et al., 2019). CTF estimation was conducted using cryoSPARC patch CTF estimation (Punjani et al., 2017). For the dataset of apo hybrid *M. tuberculosis* cytochrome *bcc*, 1,208,054 particles were picked automatically using EMD-9610 map as the template and extracted with a box size of rescaled 256 pixels (binned 2). 327,188 particles were selected after two rounds of 2D classification. 100,000 particles were used to perform Ab-Initio reconstruction in four classes, and these four classes were used as 3D volume templates for heterogeneous refinement with all selected particles. 112,804 particles converged into one class with clear signals and then re-extracted with 512 pixels. Next, this particle set was used to do homogeneous refinement and local refinement, yielding the final resolution 2.68 Å. For the dataset of Q203-bound and TB47-bound *M. tuberculosis* cytochrome *bcc*, the data processing was performed in a similar pipeline, resulting in the final reconstruction resolution at 2.67 Å and 2.93 Å, respectively (detailed parameters shown in supplementary figures).

### Model building and validation

The *M. smegmatis* respiratory complex CIII_2_CIV_2_ (PDB code: 6ADQ) model (Gong et al., 2018) as rigid body was fitted into EM density maps using UCSF Chimera 1.12 (Pettersen et al., 2004). Next, the resultant atomic model was manually modified according to the subunit sequences of *M. tuberculosis* cytochrome *bcc* and refined in COOT 0.8.9.1 (Emsley et al., 2010), followed by real-space refinement in PHENIX (Adams et al., 2010). The smile strings of Q203 and TB47 were generated and copied from ChemDraw (Li et al., 2004) and defined in PHENIX elBOW. Q203 and TB47 were manually built into the corresponding EM densities. The local resolution map was calculated in cryoSPARC (Punjani et al., 2017). All reported resolutions were based on the gold-standard FSC 0.143 criteria (Rosenthal and Henderson, 2003).

### Creation of Figures

All the figures were created using UCSF Chimera (Pettersen et al., 2004) or PyMOL (Schrodinger, 2010).

## Data availability

The accession numbers for the 3D cryo-EM density map of apo, Q203-bound and TB47-bound hybrid supercomplex in present study are EMD-30943, EMD-30944 and EMD-30945, respectively. The accession numbers for the coordinates for the apo, Q203-bound and TB47-bound hybrid supercomplex in this study are PDB: 7E1V, PDB: 7E1W and PDB: 7E1X, respectively. Correspondence and requests for materials should be addressed to the corresponding authors. The following datasets were generated:

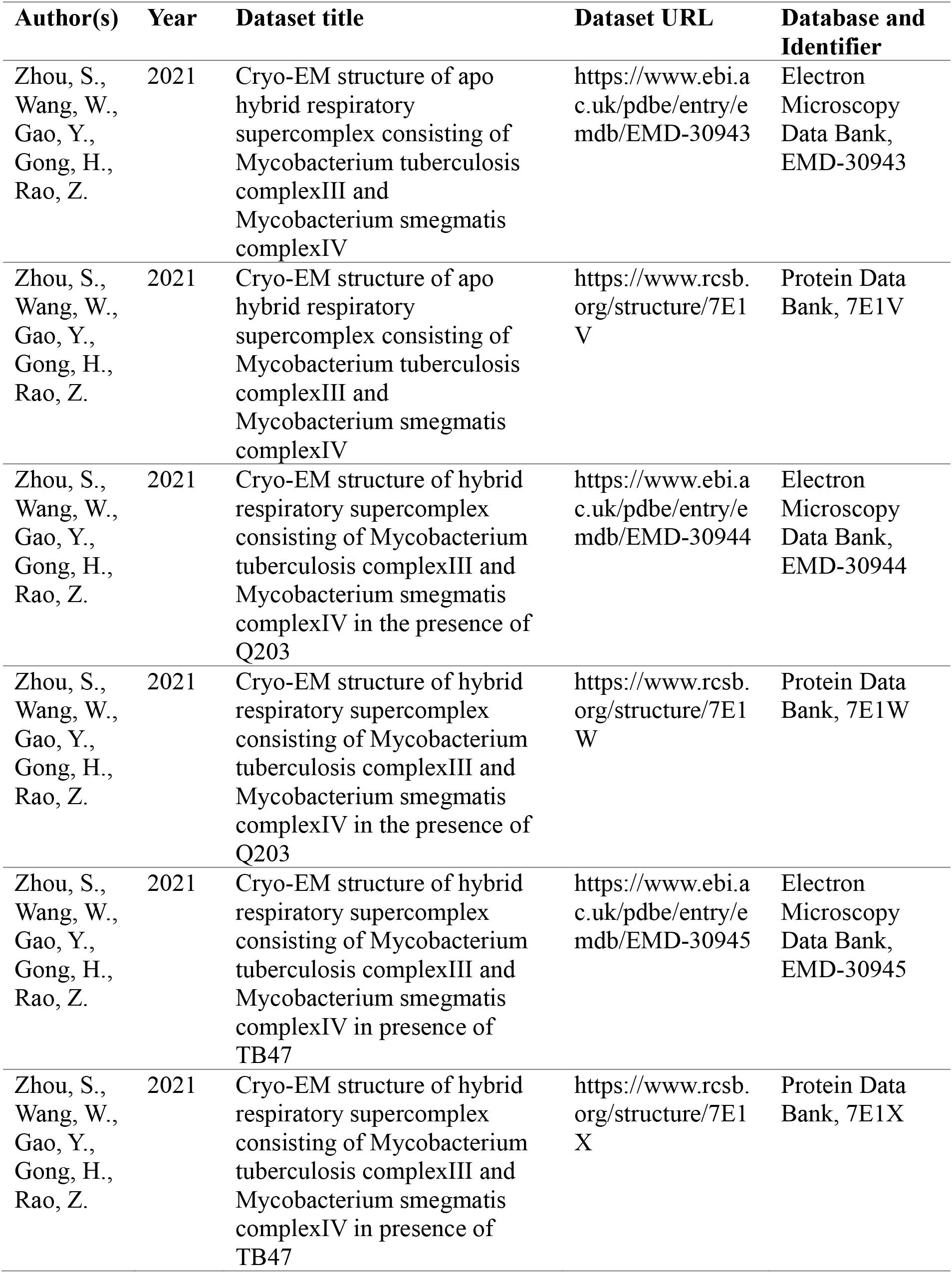

| **Author(s)** | **Year** | **Dataset title** | **Dataset URL** | **Database and Identifier** |
| --- | --- | --- | --- | --- |
| Zhou, S.,<br>Wang, W.,<br>Gao, Y.,<br>Gong, H.,<br>Rao, Z. | 2021 | Cryo-EM structure of apo hybrid respiratory supercomplex consisting of Mycobacterium tuberculosis complexIII and Mycobacterium smegmatis complexIV | https://www.ebi.ac.uk/pdbe/entry/emdb/EMD-30943 | Electron Microscopy Data Bank, EMD-30943 |
| Zhou, S.,<br>Wang, W.,<br>Gao, Y.,<br>Gong, H.,<br>Rao, Z. | 2021 | Cryo-EM structure of apo hybrid respiratory supercomplex consisting of Mycobacterium tuberculosis complexIII and Mycobacterium smegmatis complexIV | https://www.rcsb.org/structure/7E1V | Protein Data Bank, 7E1V |
| Zhou, S.,<br>Wang, W.,<br>Gao, Y.,<br>Gong, H.,<br>Rao, Z. | 2021 | Cryo-EM structure of hybrid respiratory supercomplex consisting of Mycobacterium tuberculosis complexIII and Mycobacterium smegmatis complexIV in the presence of Q203 | https://www.ebi.ac.uk/pdbe/entry/emdb/EMD-30944 | Electron Microscopy Data Bank, EMD-30944 |
| Zhou, S.,<br>Wang, W.,<br>Gao, Y.,<br>Gong, H.,<br>Rao, Z. | 2021 | Cryo-EM structure of hybrid respiratory supercomplex consisting of Mycobacterium tuberculosis complexIII and Mycobacterium smegmatis complexIV in the presence of Q203 | https://www.rcsb.org/structure/7E1W | Protein Data Bank, 7E1W |
| Zhou, S.,<br>Wang, W.,<br>Gao, Y.,<br>Gong, H.,<br>Rao, Z. | 2021 | Cryo-EM structure of hybrid respiratory supercomplex consisting of Mycobacterium tuberculosis complexIII and Mycobacterium smegmatis complexIV in presence of TB47 | https://www.ebi.ac.uk/pdbe/entry/emdb/EMD-30945 | Electron Microscopy Data Bank, EMD-30945 |
| Zhou, S.,<br>Wang, W.,<br>Gao, Y.,<br>Gong, H.,<br>Rao, Z. | 2021 | Cryo-EM structure of hybrid respiratory supercomplex consisting of Mycobacterium tuberculosis complexIII and Mycobacterium smegmatis complexIV in presence of TB47 | https://www.rcsb.org/structure/7E1X | Protein Data Bank, 7E1X |

## Acknowledgments

We thank Dr. Chao Peng of the Mass Spectrometry System at the National Facility for Protein Science in Shanghai (NFPS), Zhangjiang Lab, SARI, China for data collection and analysis and Prof. Kaixia Mi (CAS Key Laboratory of Pathogenic Microbiology and Immunology, Institute of Microbiology, CAS) for sharing the strain *M. smegmatis* mc^2^ 51. We would like to thank Prof. Gregory M. Cook (School of Biomedical Sciences, University of Otago, New Zealand) and Prof. Xiaoyun Lu (School of Pharmacy, Ji’nan University, China) for kindly providing TB47 for this study. We would also like to thank the Bio-Electron Microscopy Facility of ShanghaiTech University.

## Additional information

### Funding

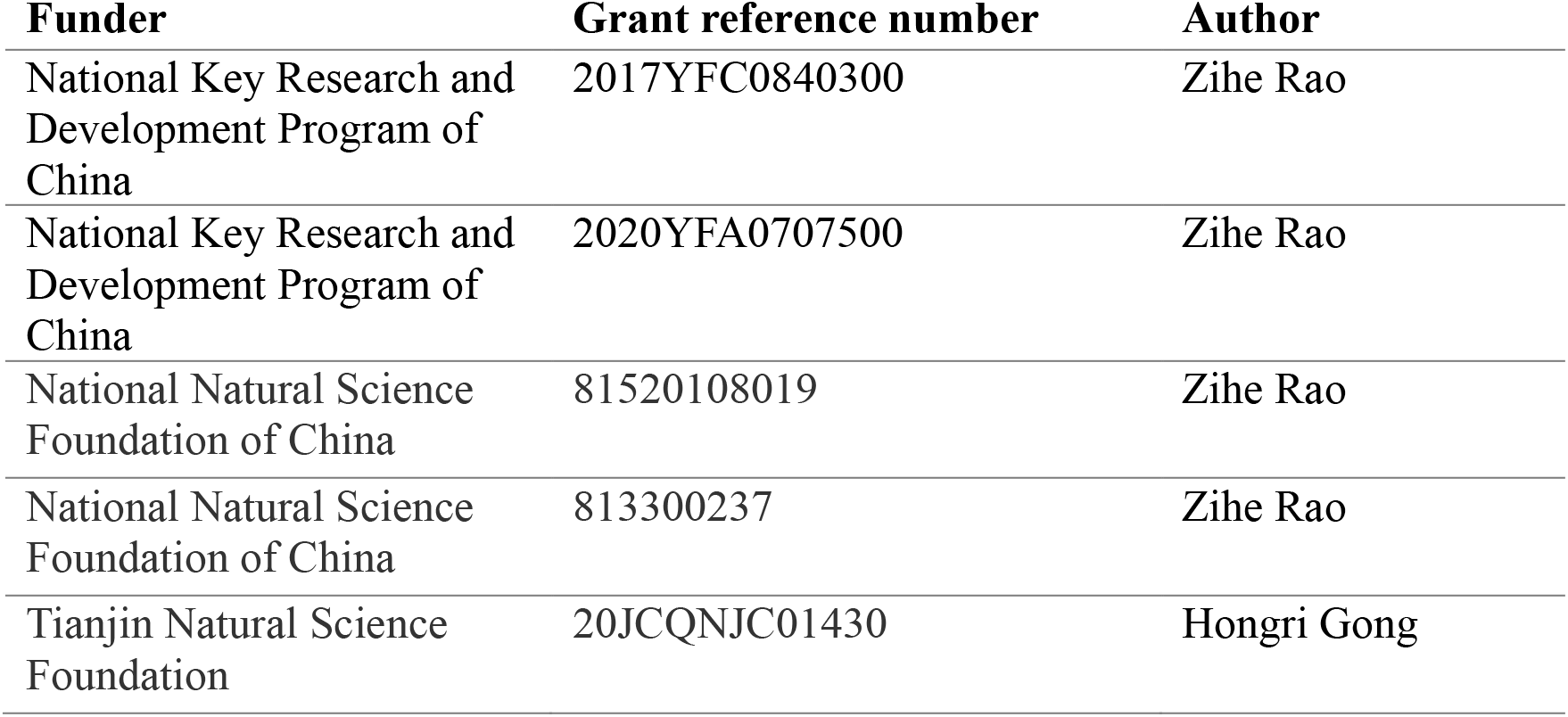

### Author contributions

Shan Zhou, Data curation, Formal analysis, Methodology, Writing—original draft, Writing—review and editing; Weiwei Wang, Data curation, Formal analysis, Methodology, Visualization, Writing— review and editing; Xiaoting Zhou, Yuying Zhang, Yuezheng Lai, Yanting Tang, Jinxu Xu, Xiaolin Yang, Quan Wang, Formal analysis, Methodology; Luke W. Guddat, Formal analysis, Writing— review and editing; Yan Gao, Data curation, Formal analysis, Methodology, Visualization, Writing—review and editing; Zihe Rao, Resources, Supervision, Funding acquisition, Writing— review and editing; Hongri Gong, Conceptualization, Investigation, Funding acquisition, Formal analysis, Methodology, Writing—original draft, Writing—review and editing.

## Tables and Figures

**Figure 2-figure supplement 1.**
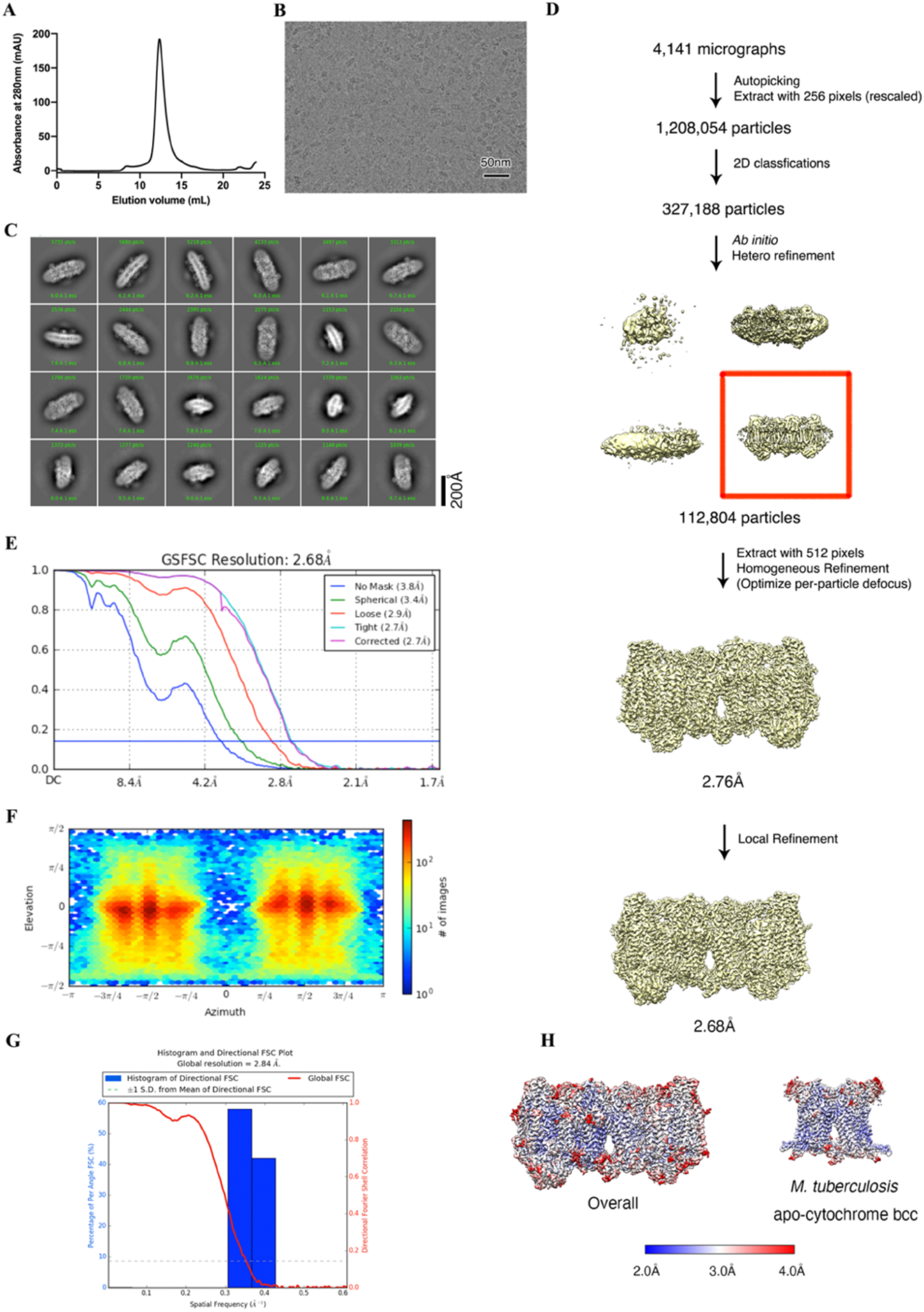
Cryo-EM data processing of apo hybrid supercomplex consisting of *Mtb* CIII and *Msm* CIV. (**A**) The elution profile of the *Mtb* cytochrome *bcc* from size exclusion chromatography (SEC). Peak fractions were pooled and concentrated for preparation on the cryo-EM grids. (**B**) Representative electron micrograph of the cryo-EM sample. (**C**)Representative 2D classification averages calculated from selected particles. (**D**) Workflow of data processing for the apo hybrid supercomplex. (**E**) FSC curves of 3D reconstructions. (**F**) Viewing direction of all particles used in the final 3D reconstruction. (**G**) 3DFSC histogram of final map. (**H**) The overall and *Mtb* cytochrome *bcc* maps, colored according to the local resolution.

**Figure 4-figure supplement 1.**
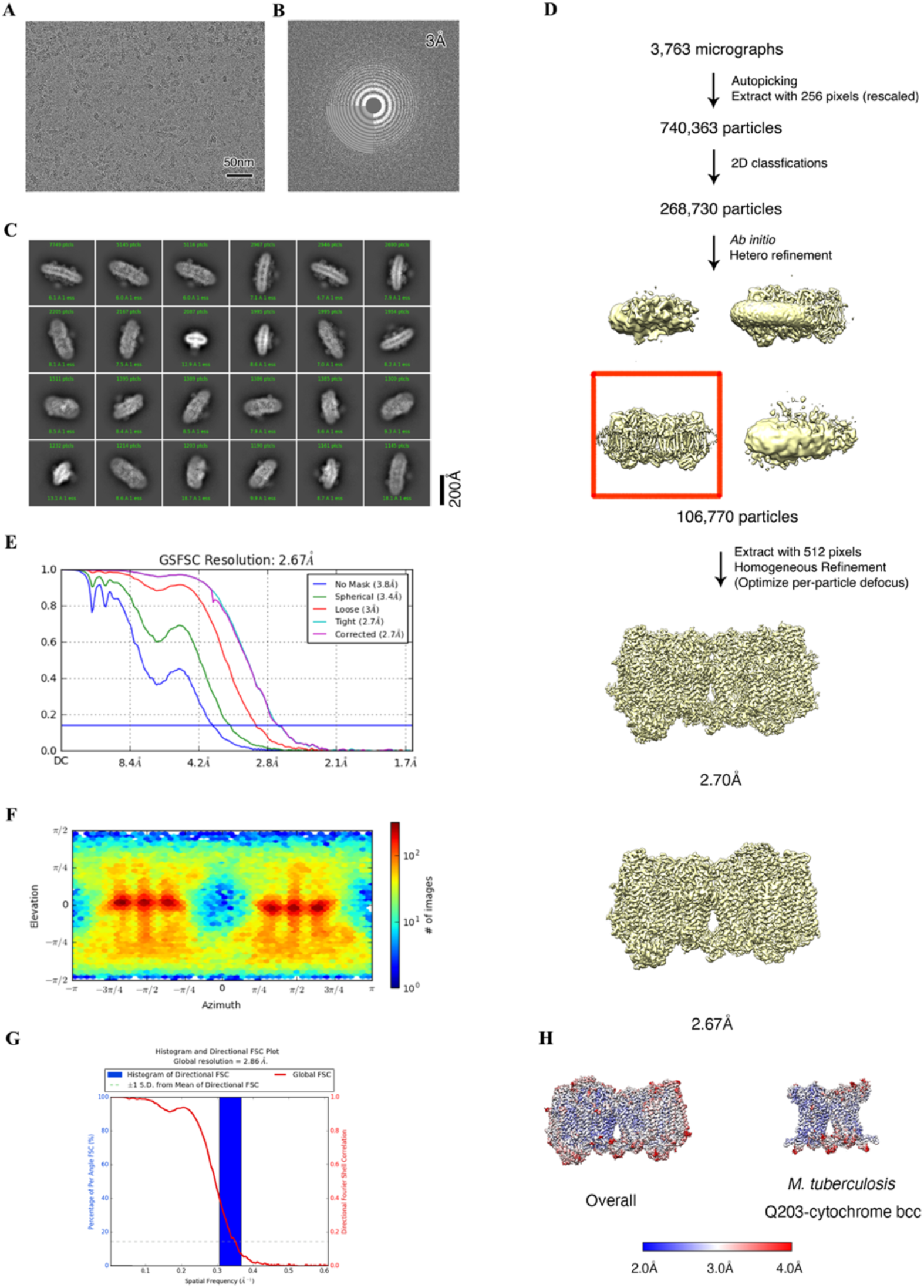
Cryo-EM data processing of hybrid supercomplex consisting of *Mtb* CIII and *Msm* CIV in the presence of Q203. (**A**) Representative electron micrograph of the cryo-EM sample. (**B**) CTF fit of motion-corrected micrographs (**C**) Representative 2D classification averages calculated from selected particles. (**D**) Workflow of data processing for the Q203-bound hybrid supercomplex. (**E**) FSC curves of 3D reconstructions. (**F**) Viewing direction of all particles used in the final 3D reconstruction. (**G**) 3DFSC histogram of final map. (**H**) The overall and *Mtb* cytochrome *bcc* maps, colored according to the local resolution.

**Figure 5-figure supplement 1.**
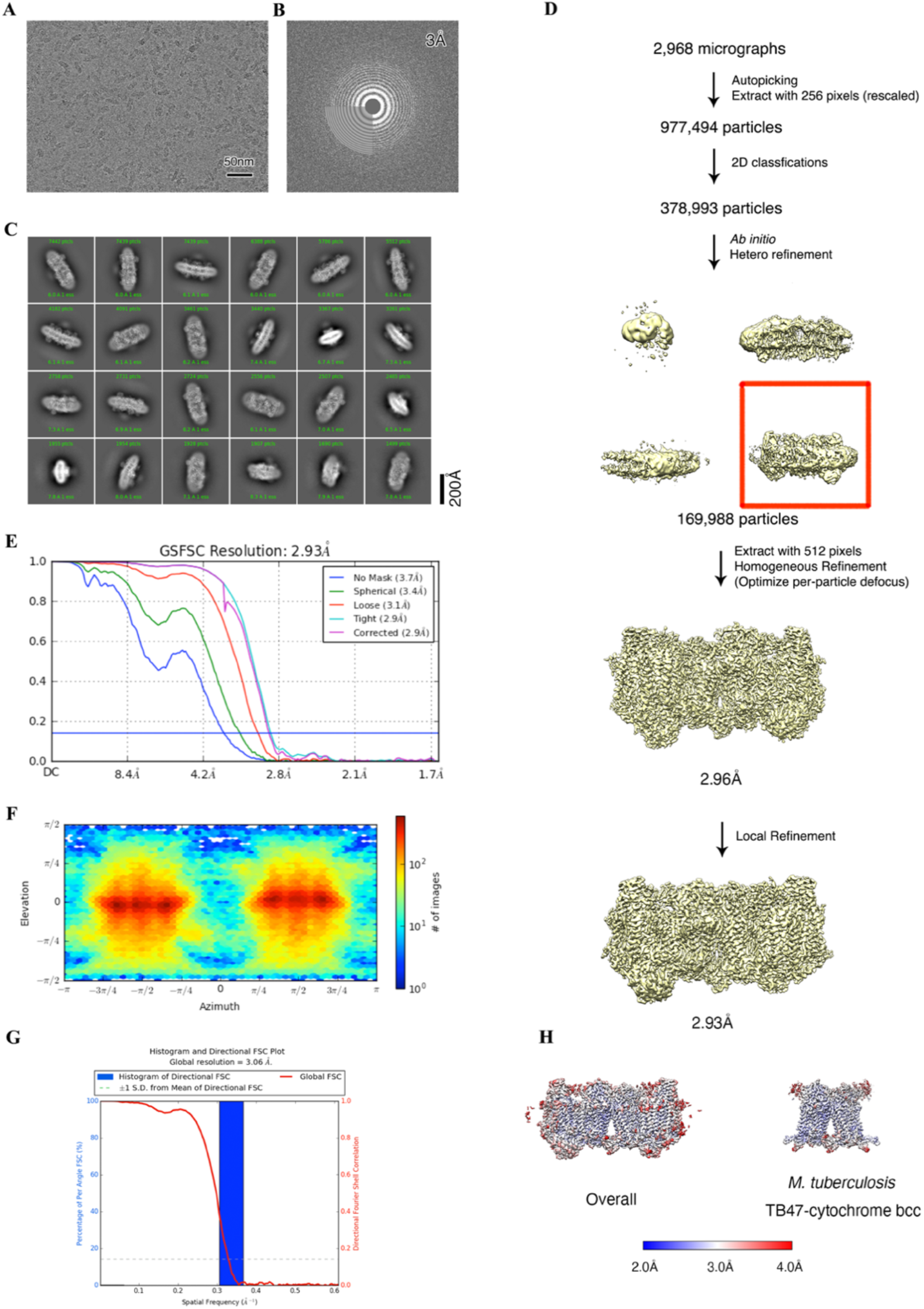
Cryo-EM data processing of the hybrid supercomplex consisting of *Mtb* CIII and *Msm* CIV in the presence of TB47. (**A**) Representative electron micrograph of the cryo-EM sample. (**B**) CTF fit of motion-corrected micrographs (**C**) Representative 2D classification averages calculated from selected particles. (**D**) Workflow of data processing for the TB47-bound hybrid supercomplex. (**E**) FSC curves of 3D reconstructions. (**F**) Viewing direction of all particles used in the final 3D reconstruction. (**G**) 3DFSC histogram of final map. (**H**) The overall and *Mtb* cytochrome *bcc* maps, colored according to the local resolution.

**Figures 2, 4, and 5-figure supplement 2.**
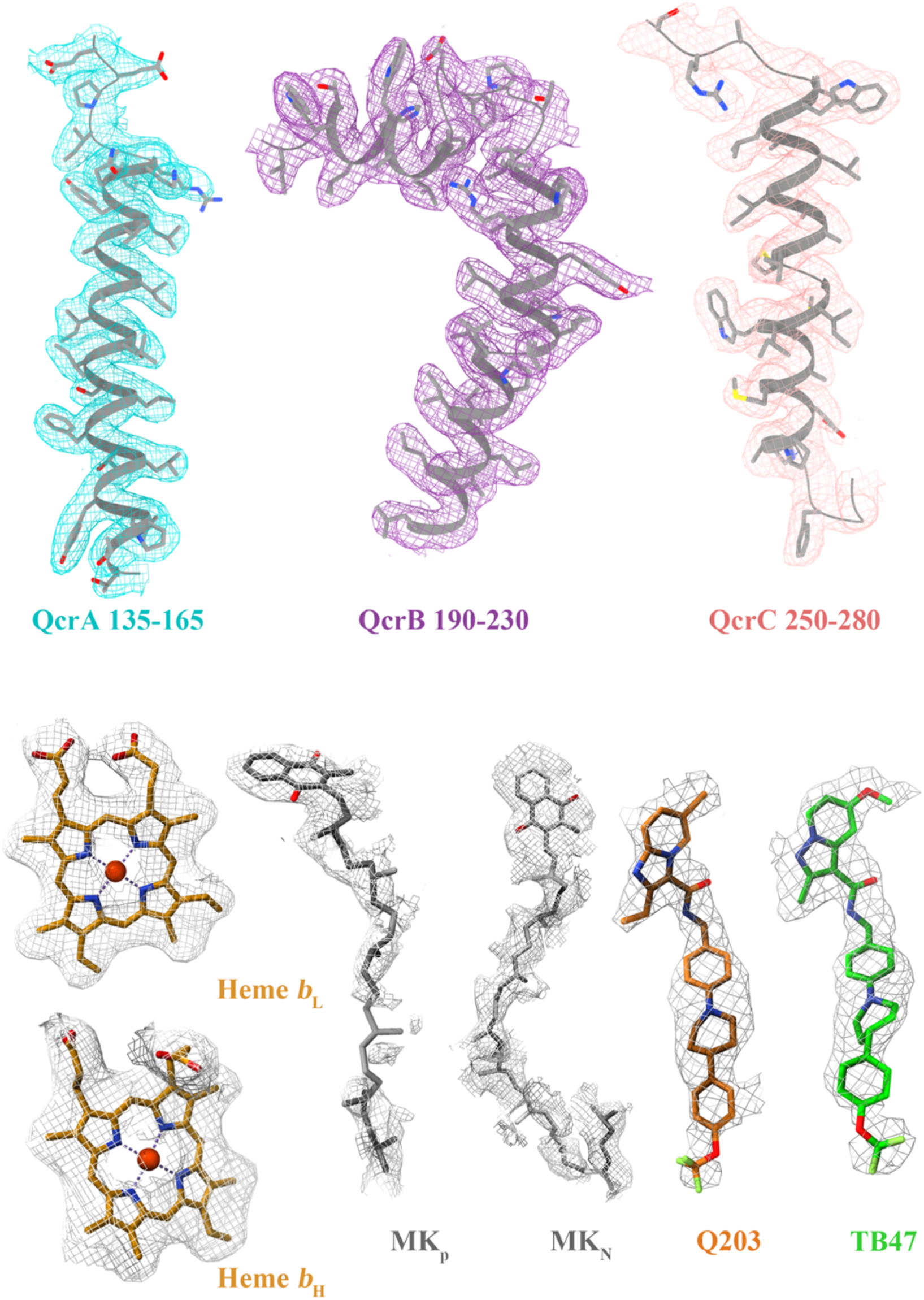
Cryo-EM map quality assessment and ligand representation of *Mtb* cytochrome *bcc* complex. Representative cryo-EM densities of individual subunits, prosthetic groups and inhibitors. Corresponding subunits with residues, prosthetic groups and inhibitors are shown in stick models or cartoon representation.

**Figures 4 and 5-figure supplement 3.**
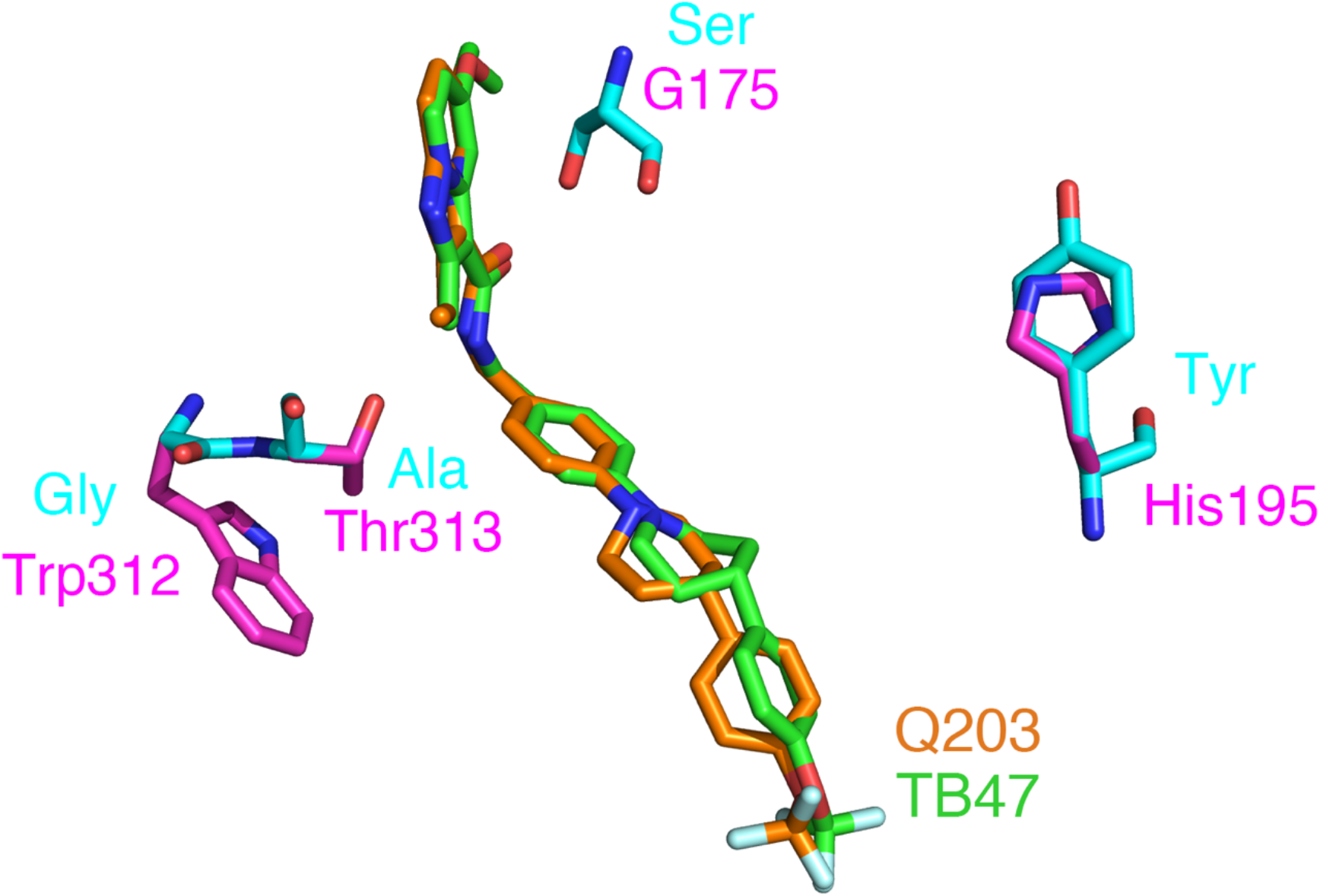
Reported mutations in Q203- and TB47-resistant *M*. ***tuberculosis*.** The native and mutant residues are colored magenta and cyan, respectively.

**Figure 6-figure supplement 1.**
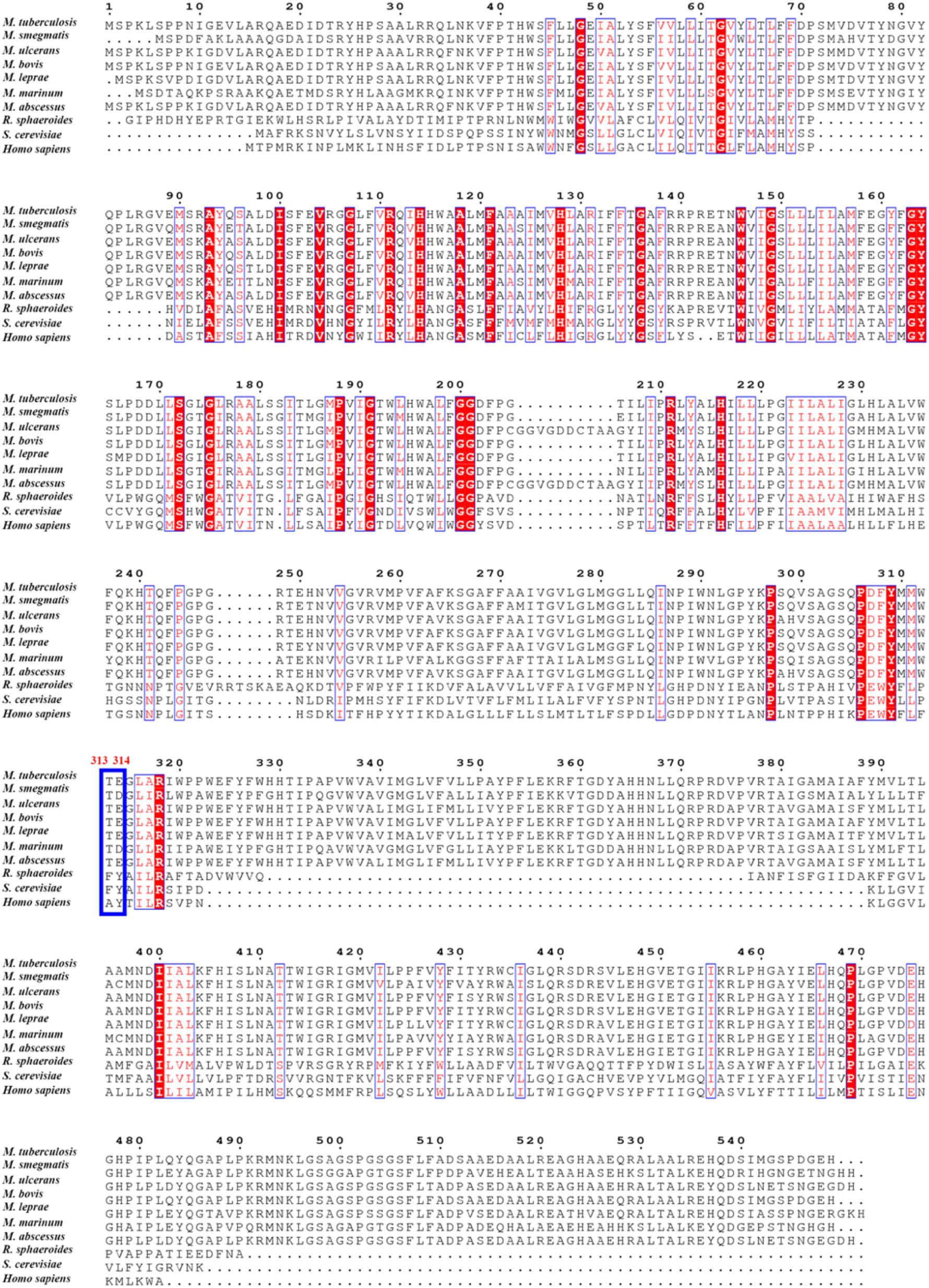
Sequence alignment of *Mtb* QcrB with the counterparts in other species and *Homo sapiens*. Residues aligned to Thr313 and Glu314 are depicted in the thick blue box. Red residues are conserved and blue indicates those less conserved.

**Figure 7-figure supplement 1.**
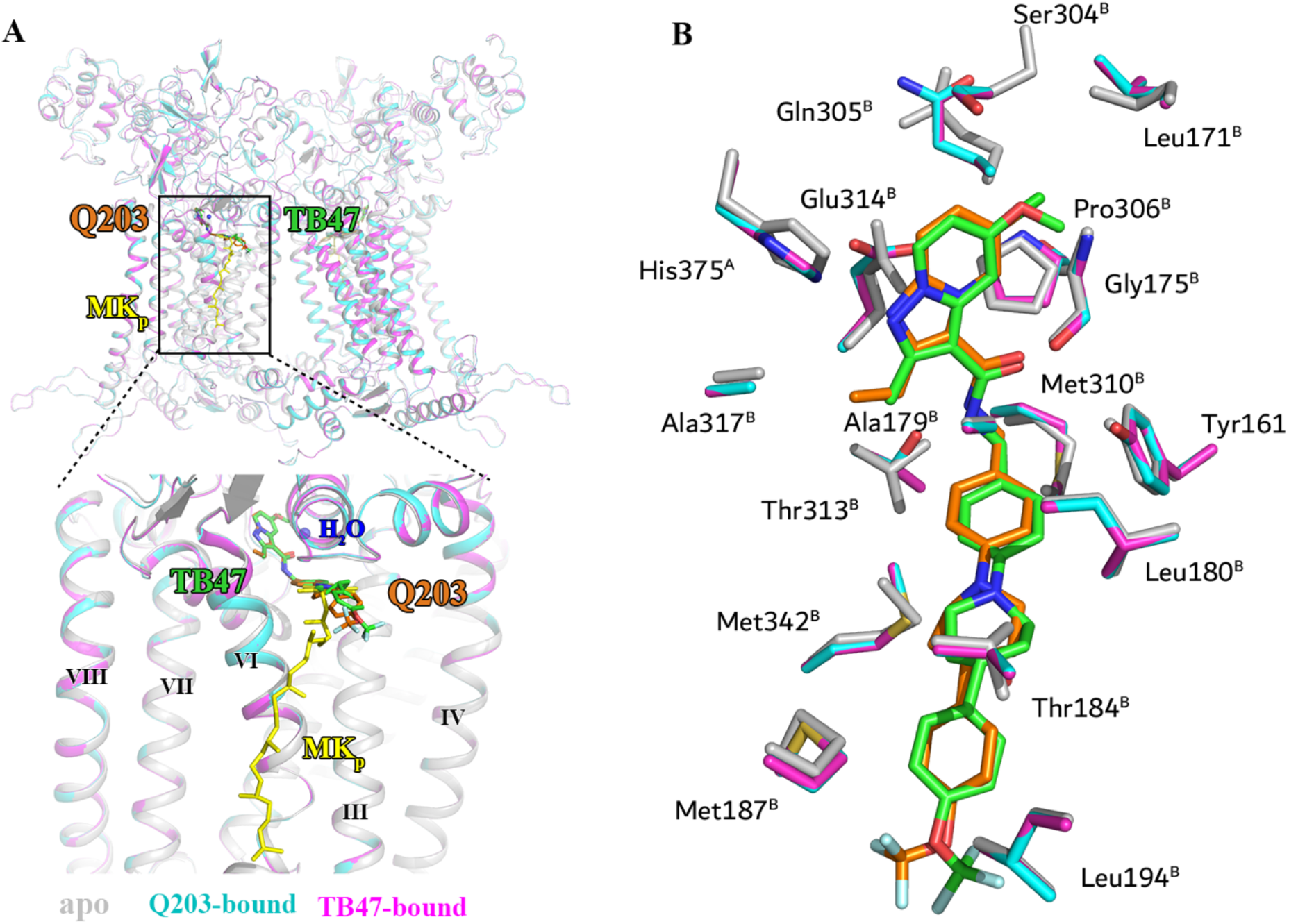
Comparison of apo and Q203/TB47-bound structures of *Mtb* cytochromes *bcc*. (**A**) Superposition of apo (gray), Q203-bound (cyan), and TB47-bound (magenta) structures of *Mtb* cytochromes *bcc*. The Q203 (orange), TB47 (green) and MK (yellow) molecules are shown as stick models, respectively. Water molecules are shown as spheres. The transmembrane helices are also labeled. (**B**) Comparison of residues surrounding Q203 (cyan sticks) and TB47 (magenta stick models) with those in apo form (gray sticks). The residues from subunits A and B are labeled with superscript A and B, respectively.

**Supplementary file 1.**
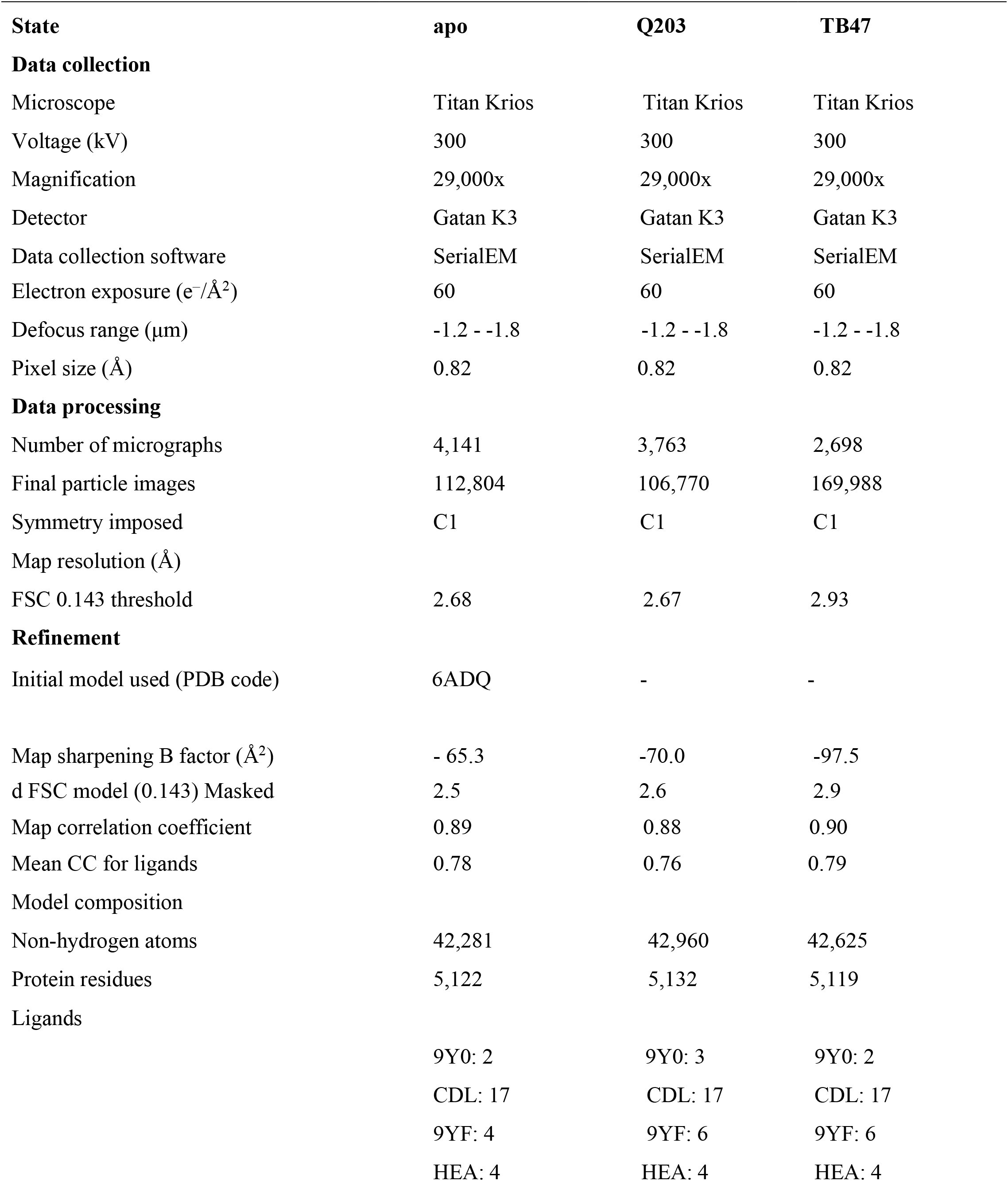
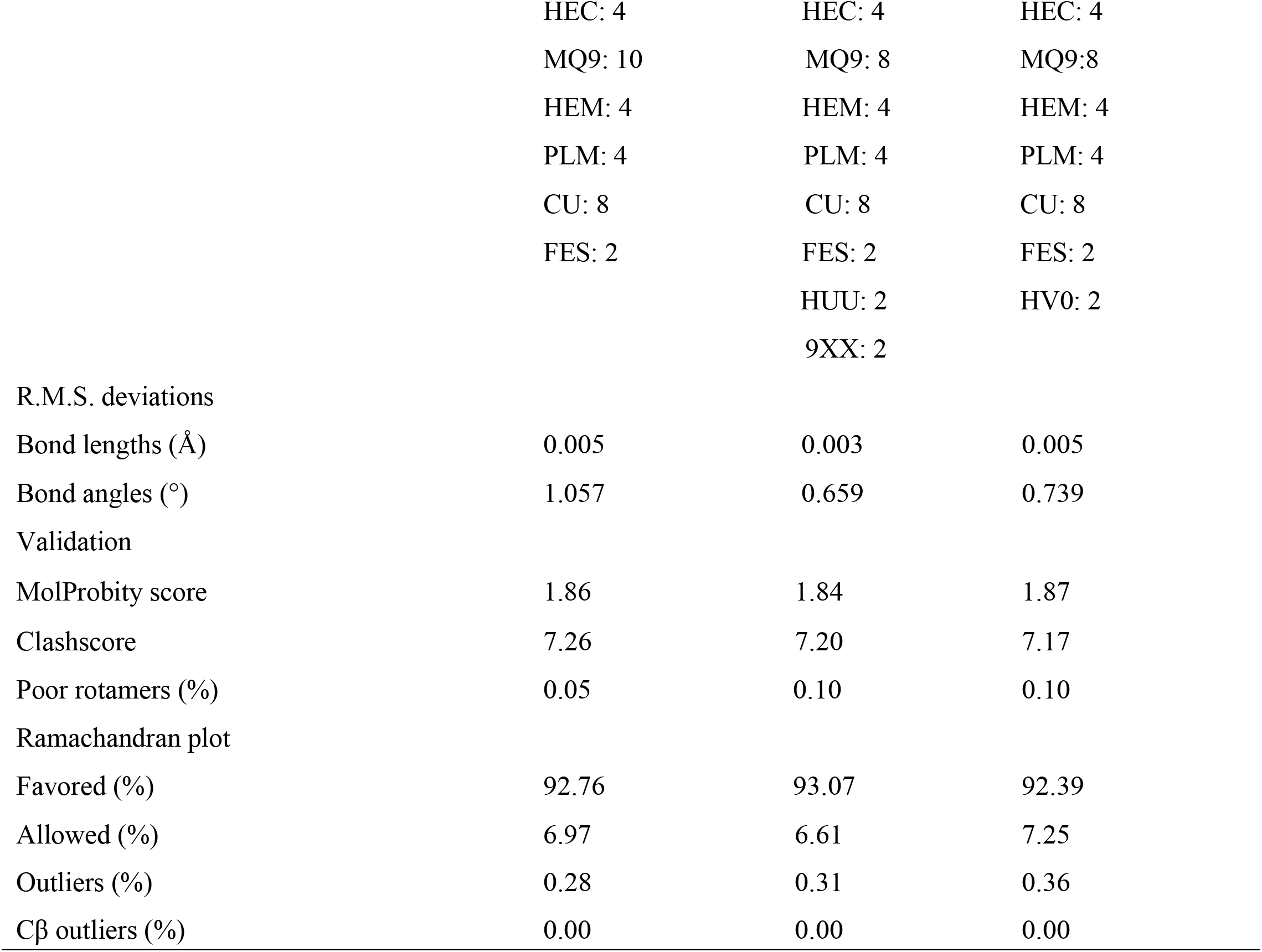
Cryo-EM data collection, refinement and validation statistics.

